# ampvis2: an R package to analyse and visualise 16S rRNA amplicon data

**DOI:** 10.1101/299537

**Authors:** Kasper S. Andersen, Rasmus H. Kirkegaard, Søren M. Karst, Mads Albertsen

**Affiliations:** Center for Microbial Communities, Aalborg University, Frederik Bajers vej 7H, 9220 Aalborg East, Denmark

## Abstract

**Summary:** Microbial community analysis using 16S rRNA gene amplicon sequencing is the backbone of many microbial ecology studies. Several approaches and pipelines exist for processing the raw data generated through DNA sequencing and convert the data into OTU-tables. Here we present ampvis2, an R package designed for analysis of microbial community data in OTU-table format with focus on simplicity, reproducibility, and sample metadata integration, with a minimal set of intuitive commands. Unique features include flexible heatmaps and simplified ordination. By generating plots using the ggplot2 package, ampvis2 produces publication-ready figures that can be easily customised. Furthermore, ampvis2 includes features for interactive visualisation, which can be convenient for larger, more complex data.

**Availability:** ampvis2 is implemented in the R statistical language and is released under the GNU A-GPL license. Documentation website and source code is maintained at: https://github.com/MadsAlbertsen/ampvis2

**Contact:** Mads Albertsen

## 1 Introduction

In the past decade, 16S rRNA gene amplicon sequencing has become the standard method for microbial community analysis (Caporaso et al., 2011; Hamady et al., 2009; Vinje et al., 2015). The approach includes several steps, from DNA extraction and barcoded PCR amplification of selected regions of the 16S rRNA gene, to sequencing and bioinformatic processing of the obtained DNA sequences (Karst et al., 2016). The latter step often involves clustering of similar sequences into Operational Taxonomic Units (OTU’s) and subsequent taxonomic assignment by comparison with a reference database of known sequences. This results in an OTU-table containing the read counts of all OTU’s in all samples as well as the assigned taxonomy. Several comprehensive pipelines exist for processing raw data and generating OTU-tables, e.g. QIIME (Caporaso et al., 2010), mothur (Schloss et al., 2009) and UPARSE (Edgar, 2013). For data analysis a popular choice is the phyloseq R package (McMurdie & Holmes, 2013), which performs many common analyses of ecological data. For the non-expert users, however, the complex S4 object system used in phyloseq to handle the data is difficult to understand and modify. Furthermore, using sample metadata, e.g. temperature, pH, sampling-site etc, to supplement the analysis remains difficult in many software tools.

Here we introduce ampvis2, an R package designed for analysis of microbial community data with a strong focus on simplicity, metadata integration, and reproducibility with a minimal set of intuitive commands. By generating plots using the ggplot2 package (Wickham, 2009), all ampvis2 plots are of publication-ready quality and easily customised by taking advantage of the vast open-source ggplot2 visualisation framework and community. Furthermore, ampvis2 aims to ease analysis of the increasingly larger data sets obtained by Next-Generation Sequencing (NGS) technologies, by enabling interactive visualisation where possible, making it possible to zoom in the plots and hover for supplemental information.

## 2 Implementation and features

The raw data files used by ampvis2 is at minimum two files; an **OTUtable** including the assigned taxonomy (from Kingdom to Species level) for each OTU, and corresponding sample **metadata**, both in a simple rowby-column structure. Optionally, the DNA sequences of the OTU’s can be loaded from a FASTA file, as well as a phylogenetic tree. Importing data directly from the QIIME (Caporaso et al., 2010), mothur (Schloss et al., 2009), and UPARSE (Edgar, 2013) pipelines is also supported. The format of the sample **metadata** is a CSV file or excel sheet containing any relevant information gathered about the individual samples; from measurements of pH, temperature or any other physical parameter, to where and when the sample was taken, content type, treatment, or anything else. The sample **metadata** is particularly important for the analysis because it contains the necessary information about how the samples can be grouped together based on these variables and is used in almost all plotting functions in ampvis2. The different types of data are then combined by the amp_load function into a single object, simplifying the process of subsetting and filtering of all the data at once using only two functions designed for the cause.

As example data included in the ampvis2 package is the microbial community composition of 658 activated sludge samples from 55 different Danish Wastewater Treatment Plants in the period 2006-2015 as determined by 16S rRNA amplicon sequencing using the Illumina platform (McIlroy et al., 2015) as well as a smaller subset hereof.

### 2.1 Flexible heatmaps

A unique feature of ampvis2 is a highly flexible heatmap, which provide a simple, but powerful overview of the microbial composition by simple color gradients as well as exact abundance values of the most abundant OTU’s. Microbial ecologists are often interested in looking at microbes at different taxonomic levels, therefore aggregating the OTU’s to any taxonomic level can be done with ease, where the combined read counts of all OTU’s assigned to a taxon of the particular level are then shown on the second axis. Furthermore, the relative abundance of each taxon can be shown as an average in a group of samples on the first axis as defined by one or more variables in the metadata.

**Fig. 1.**
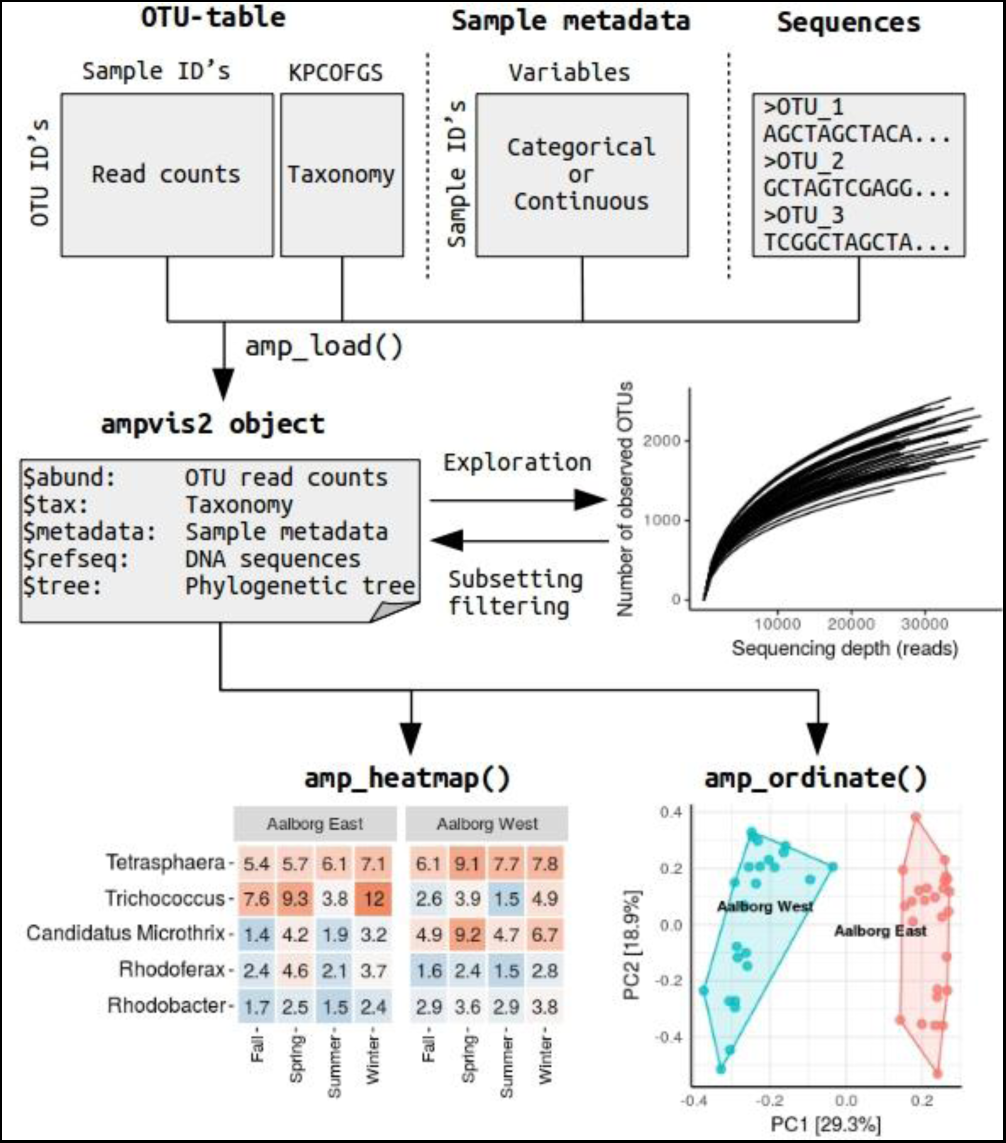
How to analyse amplicon data using ampvis2. First, the OTU-table, sample metadata, and optionally the raw DNA sequences of the OTU’s, are loaded using the amp_load() function, which checks and combines the data into a single named ampvis2 object. Secondly, the data is subject to initial exploration and quality filtering, and lastly the data can be visualised in several way using for example the amp_heatmap() or amp_ordinate() functions.

### 2.2 Simplified ordination

Due to the complex nature of compositional data, a common way to analyse and compare microbial communities is by using multivariate statistics. Perhaps one of the simplest ways to visualise the differences between samples that often contain hundreds of different OTU’s is by using ordination which seeks to reduce the dimensionality of the data and compress it into a 2-dimensional plot showing only the most important differences between the samples. The details of ordination is a subject of its own, but understanding the general principles of the most used ordination methods in microbial ecology is a useful skill as different methods can reveal different aspects of the data. Hence, ampvis2 aims to simplify the process of performing ordination, and supports seven ordination methods commonly used within the field of microbial ecology (Ramette, 2007). This is accomplished by integrating several different functions, from the vegan package (Oksanen et al., 2018) in particular, to generate advanced ordination plots with a single function call. This includes filtering of low abundant OTU’s, data transformation, calculation of a distance matrix based on common distance measures, calculation of site and species scores, fitting the correlation of environmental variable(s) in the metadata, and lastly visualisation of the result using ggplot2. The resulting plot can then be made interactive by utilizing the Plotly Javascript library (https://plot.ly).

### 2.3 Additional functionality

A number of additional functions are available within the ampvis2 package that support commonly used analysis and have detailed documentation available online. These include functions that enable effortless sub-setting, exporting, rarefaction curves, alpha-diversity calculations, boxplots, core-community analysis, time-series plots and co-occurrence network plots.

